# Elastase treatment of tendon specifically impacts the mechanical properties of the interfascicular matrix

**DOI:** 10.1101/2020.09.18.303081

**Authors:** Marta S. Godinho, Chavaunne T. Thorpe, Steve E. Greenwald, Hazel R. C. Screen

## Abstract

The tendon interfascicular matrix (IFM) binds tendon fascicles together. As a result of its low stiffness behaviour under small loads, it enables non-uniform loading and increased overall extensibility of tendon by facilitating fascicle sliding. This function is particularly important in energy storing tendons, with previous studies demonstrating enhanced extensibility, recovery and fatigue resistance in the IFM of energy storing compared to positional tendons. However, the compositional specialisations within the IFM that confer this behaviour remain to be elucidated. It is well established that the IFM is rich in elastin, therefore we sought to test the hypothesis that elastin depletion (following elastase treatment) will significantly impact IFM, but not fascicle, mechanical properties, reducing IFM resilience in all samples, but to a greater extent in younger tendons, which have a higher elastin content. Using a combination of quasi-static and fatigue testing, and optical imaging, we confirmed our hypothesis, demonstrating that elastin depletion resulted in significant decreases in IFM viscoelasticity, fatigue resistance and recoverability compared to untreated samples, with no significant changes to fascicle mechanics. Ageing had little effect on fascicle or IFM response to elastase treatment.

This study offers a first insight into the functional importance of elastin in regional specific tendon mechanics. It highlights the important contribution of elastin to IFM mechanical properties, demonstrating that maintenance of a functional elastin network within the IFM is essential to maintain IFM and thus tendon integrity.

## Introduction

Tendons possess a highly organised fibre composite structure, in which the primarily collagenous subunits are surrounded by a softer matrix material. At the highest structural level, the collagen-rich fascicles are bound together by a proteoglycan- and elastin-rich interfascicular matrix (IFM, also referred to as endotenon) [1, 2].

Whilst all tendons transfer force from muscle to bone, the stresses and strains to which different tendons are subjected vary considerably, and many highly loaded tendons additionally act as energy stores, with increased extensibility and recoverability to improve locomotion efficiency [3, 4]. Previous studies have demonstrated that the specialised mechanical properties of energy storing tendons such as the equine superficial digital flexor tendon (SDFT), primarily result from compositional and mechanical specialisation in the IFM [5–7]. The IFM enables non-uniform loading and increased overall extensibility of tendon as a result of low stiffness behaviour under small loads, which enables fascicle sliding. The capacity for fascicle sliding has been shown to be significantly greater in energy storing tendons, likely enabled by enrichment of the energy storing tendon IFM with elastin and lubricin [8, 9].

With further studies demonstrating that IFM extensibility and fatigue resistance both reduce with ageing [10, 11], it has been hypothesised that the increased injury risk seen in aged energy storing tendons originates from ageing changes in the IFM, and that poor IFM specialisation may be a primary cause of tendon overload damage. Unravelling the mechanisms that facilitate energy storage and their impact on injury risk has exciting implications for treating tendon injuries, not only providing the functional understanding from which to develop treatments, but also for targeting preventive approaches, such as mitigating the age-related loss of tendon resilience.

Recent studies have shown that, while the IFM is composed of a variety of collagens, predominantly types I and III, it is also rich in proteoglycans, particularly lubricin, and is a highly cellular region of the tendon [7, 8]. Whilst elastin only makes up approximately 5% of the dry weight of tendon, it is predominantly localised to the IFM, with light microscopy imaging suggesting elastin may bridge adjacent fascicles [8, 9]. Elastin is characterised by highly compliant and resilient behaviour, and has previously been shown to be more abundant in energy storing tendons [1, 8, 9, 12], suggesting that it may be of particular importance in tendon energy storage. Further, elastin content decreases with ageing in the energy storing SDFT, which may contribute to the increased injury risk observed with ageing specifically in energy storing tendons [9, 13, 14].

Previous studies focussed on elucidating the contribution of elastin to tendon mechanics have identified alterations in failure stress and strain [15], stiffness [16] and shear response [17] as a result of elastin depletion, utilising either heterozygous knockout models or enzymatic digestion of elastin. However, to the authors’ knowledge, no studies have investigated the effect of elastin depletion in energy storing tendons, or directly determined the effect of elastin depletion on IFM mechanics.

The current study thus focuses on a detailed exploration of the role of elastin in tendon function, carrying out a region-specific analysis of how elastin depletion impacts IFM and fascicle function in energy storing tendons, and how this is affected by ageing.

We utilise the horse superficial digital flexor tendon (SDFT) as a relevant and accepted energy storing tendon model, showing similar disease pathology and epidemiology to that seen in human energy storing tendons [14]. We combine enzymatic digestion with a series of specialised mechanical tests to test the hypothesis that elastin depletion will significantly impact the mechanical properties of the IFM, but not those of the fascicle, reducing IFM resilience in all samples, but to a greater extent in younger tendons.

## Methods

### Sample Collection and Preparation

SDFTs from 5 young (3 to 7 years - young group) and 5 old (15 to 19 years - old group) horses were dissected from both forelimbs of horses euthanised at a commercial abattoir within 24 hours post-mortem. Tendons were divided into four equal sized longitudinal sections, wrapped in phosphate-buffered saline (PBS) dampened tissue and aluminium foil, and stored at −20°C until required, enabling the multiple quasi-static, fatigue and recovery experiments to be conducted on separate sections, avoiding multiple freeze-thaw cycles.

### Optimisation and validation of elastase digestion protocol

20 samples, each composed of 2 fascicles bound together by IFM (approximately 40mm long and 1mm in diameter), were dissected from a single young SDFT section (n=1; 7 years old) to optimise and validate the elastase digestion protocol. Samples were divided into 4 groups: fresh, control, 0.2U/ml elastase and 2U/ml elastase, with the chosen enzyme concentrations based on previous literature [18] in conjunction with a preliminary study. Samples in the “fresh” group were stored at 4 °C and analysed within 16 hours of dissection, whilst samples in the two elastase groups and the control group were incubated in a buffer solution with and without the inclusion of elastase for 16 hours at room temperature, with gentle agitation. The buffer solution comprised 5 ml of 1x PBS plus 0.1mg/ml soybean trypsin inhibitor (SBTI) solution. Elastase (trypsin-free porcine pancreatic elastase, EPC134, Elastin Products Co., Owensville, MO) was added at concentrations of 0.2U/ml or 2U/ml, to the desired groups.

After incubation, samples were washed in PBS and divided into 2 groups for biochemical and immunohistochemical analyses. Samples for biochemical analysis were stored at −20°C until required, while samples for immunohistochemical analysis were prepared immediately.

#### Elastin immunolocalisation

One sample from each test group was immunolabelled for elastin and cell nuclei. Samples were fixed in 4% paraformaldehyde (PFA) for 30 minutes, washed in PBS, then incubated in 10% Goat Serum for 1H, followed by the elastin antibody (Ab9519; 1:100 dilution in 5% goat serum) overnight. Samples were then washed in PBS, incubated in the secondary antibody (555 Goat anti Mouse IgG H+L, 1:500 in 5% goat serum) for 1H, washed again in PBS, and finally incubated for a further 5min in DAPI (1:1000 in 5% goat serum).

Samples were placed on poly-lysine slides, mounted with prolong Diamond antifade and imaged with a laser scanning confocal microscope (Zeiss ELYRA; Carl Zeiss AG, Oberkochen, Germany) using a 63x oil objective. Confocal z series were taken with an image size of 225 x 225μm, pixel size of 0.11 x 0.11μm and a z-step size of 0.25μm.

#### Biochemical Analysis

The amounts of elastin, sulphated glycosaminoglycan (GAG) and collagen type I, were quantified in fresh, control, 0.2U/ml and 2U/ml elastase treatment groups, combining the remaining 4 samples in each treatment group to ensure ~25mg dry weight of tissue, on which to perform all three biochemical assays.

Samples were powdered using a Micro dismembrator, freeze dried and then weighed. Approximately 6mg of powered sample was used to determine elastin content with the Fastin Elastin assay (Biocolor, UK). To briefly describe the procedure (fully detailed in [9]), elastin was extracted from the tendon samples using oxalic acid, and alpha-elastin was used to create a standard curve.

The remaining tissue (10-20mg dry weight) was solubilised in papain (P3125, Sigma, UK) for 18h at 60°C, prior to measurement of sulphated GAG and collagen content using standard DMMB and hydroxyproline assays respectively [19, 20].

Results revealed that treatment with 2U/ml elastase caused a 70% reduction in elastin content, and therefore this concentration was selected for all subsequent experiments.

### The effect of elastin depletion on fascicle and IFM mechanical properties

Approximately 45 fascicles and 45 IFM samples (approx. 40 mm in length) were dissected from the mid-metacarpal region of each SDFT quarter as described previously [21, 22], and divided into 3 groups: fresh, control and 2U/ml elastase (n = 15 samples per group from each biological replicate). Samples in the “fresh” group were maintained for up to 16 hours in wet tissue paper at 4°C until testing, whilst samples in the elastase and control groups were incubated in buffer solution under gentle agitation, with and without the inclusion of elastase, for 16 hours at room temperature prior to the start of testing. After incubation, samples were rinsed twice in PBS then maintained on tissue paper dampened with PBS, ready for testing. For each sample, fascicle diameter was first measured using a non-contact laser micrometer [21, 22], assuming a circular shape to calculate cross section area (CSA). The mechanical properties were then determined using an electrodynamic testing machine (Instron ElectroPuls 1000) with a 250N load cell, as previously described [5]. Briefly, fascicles were secured in pneumatic grips (grip to grip distance: 20mm; gripping pressure 4 bar) and pre-loaded to 0.1N, which represents approximately 2% of fascicle typical failure load. Fascicles were then preconditioned with 10 sinusoidal loading cycles between 0 and 3% strain (approx. 25% of failure strain; frequency: 1Hz), immediately followed by a pull to failure test at a strain rate of 5% per second.

IFM samples were secured in the same manner and pre-loaded to the smallest positive load value that could be detected (approximately 0.02N). Samples were pre-conditioned with 10 loading cycles between 0 and 0.5mm of extension (approx. 25% of failure extension; sine wave; frequency: 1Hz), and pulled apart to failure at a speed of 1mm/s.

All samples were kept hydrated with a mist of PBS during testing. Force and displacement data for all samples were continuously recorded at 100 Hz during both preconditioning and pull to failure, and where appropriate, engineering stress and strain were calculated using the CSA and effective gauge length, respectively. Force data were smoothed, prior to any calculations, using a 9-point moving average filter, to remove noise [21]. Displacement at which the initial pre-load was reached was taken as the start point for the test to failure in both fascicles and IFM samples. Maximum stiffness (for IFM samples) or modulus (for fascicles) was identified by taking continuous tangent calculations across every 9 data points of the respective pull to failure curve, then identifying the peak value. Hysteresis was calculated as the difference in area under the loading and unloading curves from the first to tenth preconditioning cycles. After analysis, any fascicle or IFM sample in which failure properties or maximum modulus/stiffness were more than 2.5 times above or below the standard deviation of the mean, were excluded.

### The effect of elastin depletion on IFM fatigue properties

To investigate the effects of elastase treatment on IFM fatigue properties, the fatigue properties of fresh, control and elastase treated IFM samples (n=15 samples per group from each biological replicate) were explored, using a mechanical testing machine (Electroforce 5500, TA instruments, Delaware, USA), housed within a cell culture incubator (37 °C, 20% O_2_, 5% CO_2_), with a 22N load cell. Samples were dissected as previously described, then secured in a custom designed chamber (grip to grip distance of 10mm) which was filled with PBS to prevent samples from drying out. IFM samples were pre-loaded to 0.02N and the load for creep tests was determined by first carrying out a single displacement-controlled cycle to 1 mm extension, and selecting the peak load reached. We have previously shown this to equate to approximately 50% of the predicted failure extension and to give the most consistent conditions for a controlled creep test [23]. The identified load was applied cyclically to the samples at a frequency of 1 Hz until sample failure, with maximum and minimum displacement data continuously recorded at 100 Hz throughout the tests.

From the resulting creep curves, the number of cycles to failure, the creep between cycles 1 and 10 (mm), and the secondary creep rate (mm/cycle) were calculated. Data were compared between treatments and age groups. Samples in which no secondary creep was evident (immediate failure) were assumed to be damaged and rejected from the data set.

### The effect of elastin depletion on IFM recovery

A custom designed tensile straining rig was used to investigate the ability of the IFM to recover from loading in fresh, control and elastase treated samples (n = 2 samples per group from each biological replicate). IFM samples were prepared with a 10mm test region, by cutting opposing ends from two adherent fascicles as previously described, and secured in the rig at a 15mm grip-to-grip distance, such that a single intact fascicle was held at each end, with the 10mm IFM testing region in the middle (Figure 1a). Once samples were secured, four equally spaced lines were manually drawn across the 10mm IFM testing region using a permanent marker pen. Grips were then secured into the rig and the sample immersed in PBS. A Canon EOS 700D camera with a Sigma 105mm F2.8 EX DG MACRO OS, fixed to a tripod was placed directly above the sample at a consistent height and location, to allow visualisation of sample movement. Controlled by linear actuators, the grips were slowly moved apart (at 0.05mm/s), applying small increments of displacement, whilst visually monitoring the sample until it lifted slightly off the base of the rig, which provided a consistent start point for tests [24].

**Figure 1.**
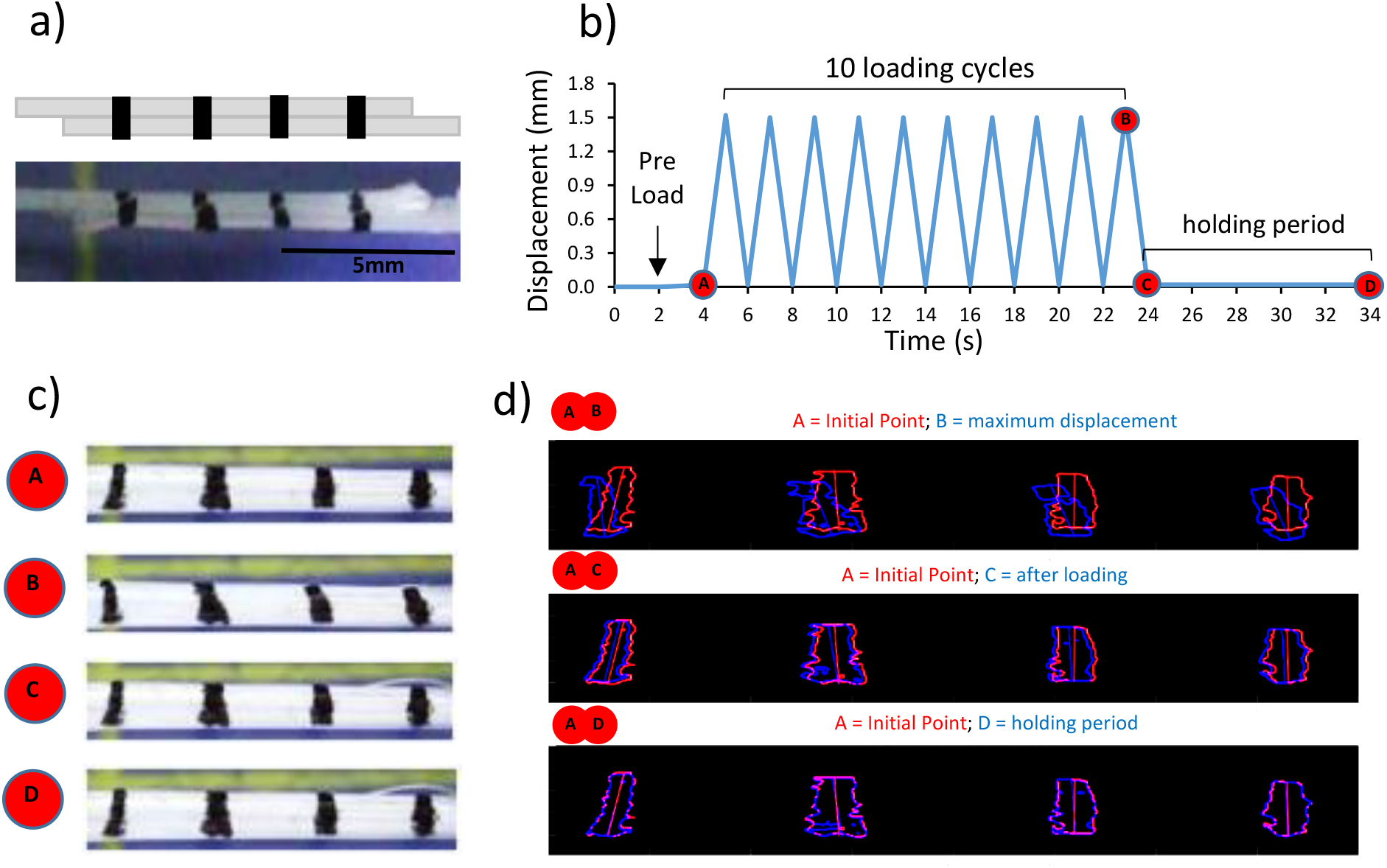
Schematic showing IFM recovery testing protocol. To test the ability of the IFM to recover, the opposite ends of each fascicle were removed, so that only 10mm of intact IFM was left connecting the fascicles. Four lines were drawn across the two fascicles in the central test region to track local displacements; samples shown schematically and pictorially, where samples are pictured on a blue cutting board (a). A time-displacement graph pictorially represents the test protocol and times at which images were analysed. Four specific test points were identified; A: initial point, B: maximum displacement, C: immediately after removal of load and D: after a 10 second recovery period (b). Still frames of the sample were extracted from the video at each selected time point (c). Tracking algorithms were adopted to investigate the displacement of the lines, and thus fascicles, during each test (d).

Samples were subjected to 10 loading cycles at 0.5Hz between 0mm and 1.5mm (which corresponds to approximately 75% of IFM predicted failure extension), followed by a ten seconds hold period at 0mm displacement to allow for any IFM recovery (Figure 1b). Video footage was recorded at a rate of 10 frames per second throughout the test. At the end of the test, samples were pulled to failure at a rate of 1mm/s, to ensure samples had been prepared correctly with both fascicles fully cut and no intact fibres traversing the test region. Samples subsequently shown to have intact fibres traversing the testing region were excluded from the data set.

### Data Analysis

The frames relating to specific time points during the test, designated A, B, C and D (Figure 1b) were selected for further analysis, measuring the angular deviation of lines during the test (Figure 1c). To briefly describe the process, Fiji (ImageJ) was used to draw a region of interest (ROI) around the marker lines, allowing the relevant image region to be cropped (Figure 1c). The cropped images were first smoothed in MATLAB using a Gaussian filter and then thresholded, using the same parameters for all images (sensitivity 0.1; marker margin 10, chosen from preliminary experiments). Each marker line in the cropped image, was divided into a stack of horizontal lines, and for each one, the midpoint was found, and an interpolated line drawn through the midpoints (to give a line of single pixel thickness tracing the middle of the marker). The angular orientation of each line constructed in this way, was calculated relative to its orientation in the reference image (initial point - “A “), and then the average angular deviation across all four lines calculated and reported at each time point. From these data, percentage recovery after loading (comparing point C and B) and the total recovery (comparing point D and B) were also calculated.

In order to determine the potential error in measurements, the impact of shifting one end of the interpolated line by 1 pixel was investigated, demonstrating that this would affect the calculated angle by approximately 0.2 degrees. Thus, all angular deviation values lower than 0.2 degrees were excluded from the data set.

### Statistical Analysis

All statistical analyses were carried out using Minitab 17. Data were tested for normality using the Anderson – Darling test. If normally distributed, a two-way ANOVA (grouping information using the Tukey Method and 95% confidence) was used to evaluate differences between treatments and age groups. Treatment and horse age were used as factors for the ANOVA, and each donor was nested with horse age to account for the use of multiple samples from individual donors. Data that did not follow a normal distribution were first transformed using a Box Cox transformation, and if still not normally distributed, a nonparametric Mann–Whitney test was used. Results were considered statistically significant if p<0.05. Data in bar graphs are displayed as mean ± standard deviation. Box plots graphs show all data points.

## Results

### Elastin Depletion Validation

Immunolabelling confirmed the presence of elastin fibres, predominantly localised to the IFM region of both fresh (Figure 2a) and control (Figure 2b) samples. Very few elastin fibres where seen in the samples treated with 0.2U/ml elastase (Figure 2c) and no elastin fibres were seen in the 2U/ml elastase treated samples (Figure 2d). Results from biochemical analysis showed an elastin reduction of over 70% when samples were incubated in a 2U/ml elastase solution (Figure 2e). The low amount of elastin remaining might be in the form of elastin fragments trapped in the tissue, which were either too small to visualise with immunohistochemistry or digested in such a manner that the elastin antibody no longer recognised them.

**Figure 2.**
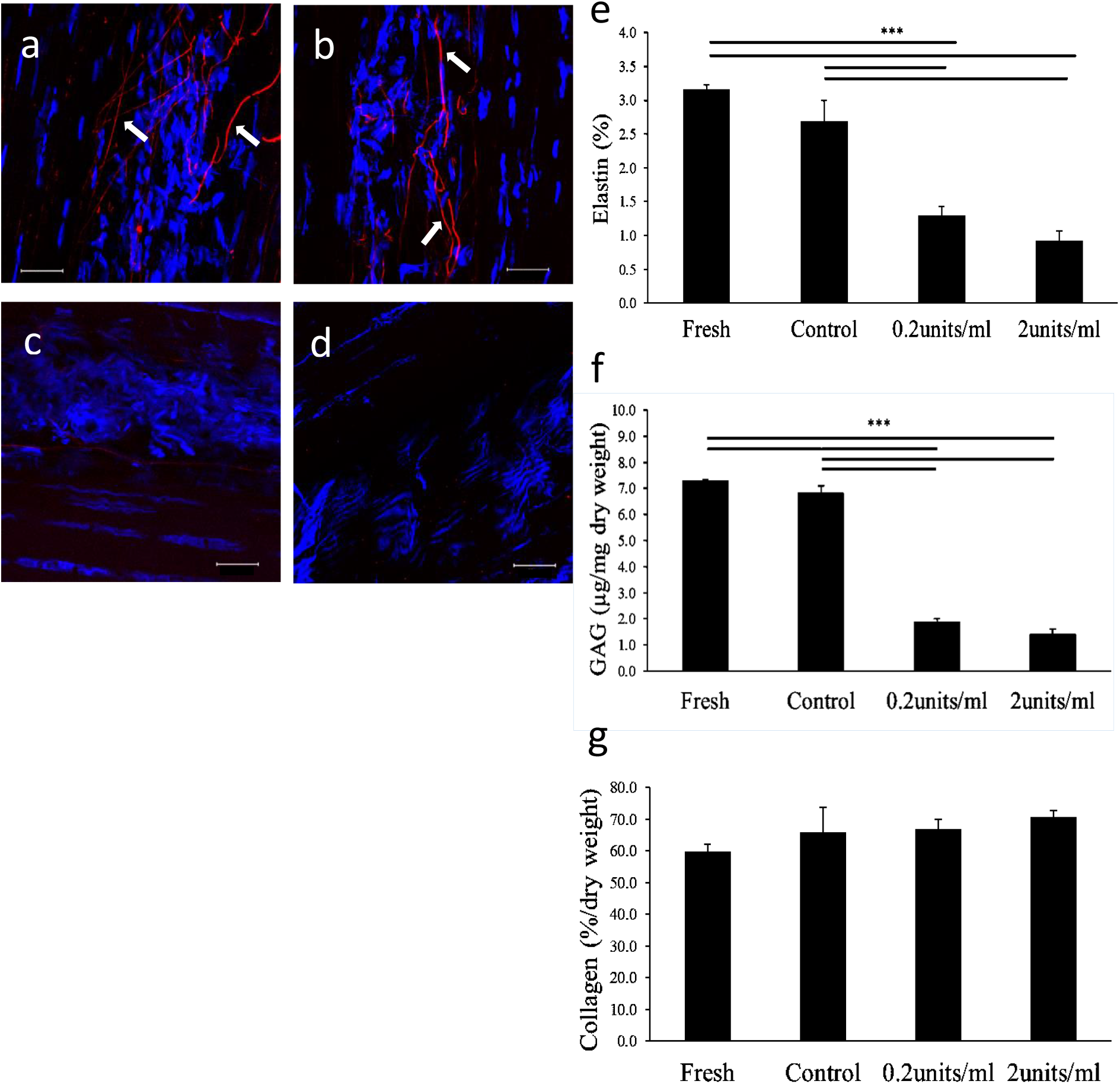
Validation of elastase treatment. (a-d) Representative confocal images showing tendon explants immunolabelled for elastin (red) and cell nuclei (blue): fresh (a), after incubation in control buffer (b), in 0.2U/ml elastase solution (c) and 2U/ml elastase solution (d). Scale bar: 20μm. Visible elastin fibres are noted with arrows. Quantitative investigation of tendon matrix composition compares elastin (e), GAG (f) and collagen (g) content for fresh, control and elastase treated samples. Significant differences between treatments are identified by: *** p<0.001 (normally distributed data – ANOVA). Data are displayed as mean ± standard deviation.

Elastase treatment also resulted in a significant decrease in GAG content, particularly in a 2U/ml elastase solution (>80%) (Figure 2f). As such a reduction in GAG content may have impacted the study findings, an additional investigation was carried out to determine the impact of GAG removal on IFM and fascicle mechanics, using chondroitinase ABC treatment for GAG removal. Full methods and results of the fascicle and IFM pull to failure studies after chondroitinase treatment are provided in the supplementary information. No differences in the mechanical properties of fascicles or IFM were evident in any of the GAG depleted samples.

Collagen content was unaffected by elastase treatment (Fig. 2g).

Based on these results, 2U/ml elastase was used in all subsequent experiments.

### Fascicle & IFM Failure Properties

Fascicle and IFM failure properties are shown in Table 1. Fascicle failure tests showed no significant differences in any mechanical parameters between treatment groups, nor any significant differences with ageing. By contrast, elastase treatment led to a significant reduction in IFM failure load and maximum stiffness in both young and old groups, and an overall increase in IFM hysteresis in elastase treated samples (Table 1). No significant differences between fresh and control samples were found in any of the parameters assessed, indicating that differences resulted from elastase treatment and not incubation. The majority of parameters were unaffected by ageing, with the exception of failure extension and initial hysteresis, which showed small but significant decreases with ageing in elastase treated IFM samples.

**Table 1.** Fascicle & IFM Failure Properties in fresh, control, and elastase treated samples from young and old horses. n=5/age group; total samples tested: 15/treatment/tendon. Data are shown as mean ± SD. Significant differences are flagged with: *p<0.05, **p<0.01 and ***p<0.001. CE indicates differences between control and elastase treated groups; FE, differences between fresh and elastase groups.

| Fascicles | Young SDFT |  |  |  | Old SDFT |  |  |  | Between age groups comparisons |
| --- | --- | --- | --- | --- | --- | --- | --- | --- | --- |
|  | Fresh (F) | Control (C) | Elastase (E) | Within Age group comparisons | Fresh (F) | Control (C) | Elastase (E) | Within Age group comparisons |  |
| Diameter (mm) | 0.24<br>±0.04 | 0.23<br>±0.04 | 0.23<br>±0.04 | | 0.23<br>±0.05 | 0.21<br>±0.05 | 0.22<br>±0.05 | | |
| Failure Load (N) | 2.16<br>±0.72 | 2.19<br>±0.89 | 2.06<br>±0.81 | | 1.99<br>±0.97 | 1.78<br>±0.92 | 1.79<br>±0.84 | | |
| Failure Stress (MPa) | 49.95<br>±14.39 | 50.69<br>±14.23 | 49.17<br>±18.78 | | 47.21<br>±18.24 | 55.23<br>±25.17 | 51.16<br>±21.44 | | |
| Strain at Failure (%) | 11.38<br>±2.41 | 11.58<br>±1.89 | 12.00<br>±2.25 | | 10.54<br>±3.01 | 10.68<br>±2.93 | 11.44<br>±2.35 | | |
| Maximum Modulus (MPa) | 649.17<br>±139.68 | 643.77<br>±149.75 | 630.23<br>±205.24 | | 631.14<br>±156.29 | 747.95<br>±260.10 | 675.69<br>±219.76 | | |
| Hysteresis Cycle 1 (%) | 25.55<br>±3.76 | 23.83<br>±3.61 | 25.65<br>±2.73 | | 23.48<br>±4.54 | 22.76<br>±4.03 | 24.13<br>±4.25 | | |
| Hysteresis Cycle 10 (%) | 9.54<br>±1.59 | 8.99<br>±1.75 | 9.28<br>±1.78 | | 8.76<br>±1.93 | 8.54<br>±1.98 | 8.86<br>±2.19 | | |
| IFM Samples | Young SDFT |  |  |  | Old SDFT |  |  |  | Between age groups comparisons |
|  | Fresh (F) | Control (C) | Elastase (E) | Within Age group comparisons | Fresh (F) | Control (C) | Elastase (E) | Within Age group comparisons |  |
| Failure Load (N) | 1.66<br>±0.69 | 1.47<br>±0.56 | 0.95<br>±0.39 | ***FE; ***CE | 1.63<br>±0.60 | 1.49<br>±0.59 | 0.86<br>±0.37 | ***FE; ***CE | |
| Failure Extension (mm) | 1.99<br>±0.55 | 2.09<br>±0.60 | 1.87<br>±0.59 | | 1.73<br>±0.48 | 1.75<br>±0.45 | 1.48<br>±0.45 | | *C; **E |
| Maximum Stiffness (N/mm) | 1.30<br>±0.36 | 1.18<br>±0.29 | 0.91<br>±0.23 | ***FE; **CE | 1.46<br>±0.41 | 1.28<br>±0.32 | 0.91<br>±0.24 | ***FE; ***CE | |
| Hysteresis Cycle 1 (%) | 31.57<br>±5.53 | 31.57<br>±3.80 | 39.20<br>±7.17 | **FE; **CE | 30.64<br>±5.67 | 33.08<br>±7.06 | 35.29<br>±6.49 | ***FE | *E |
| Hysteresis Cycle 10 (%) | 12.63<br>±3.41 | 13.05<br>±2.42 | 17.28<br>±6.63 | ***FE; **CE | 14.47<br>±4.82 | 16.11<br>±6.97 | 18.44<br>±12.55 | | |

### IFM Fatigue Properties

Representative creep curves for young and old SDFT IFM samples are shown in Figure 3, whilst all SDFT IFM fatigue data are summarised in Figure 4a-c.

**Figure 3.**
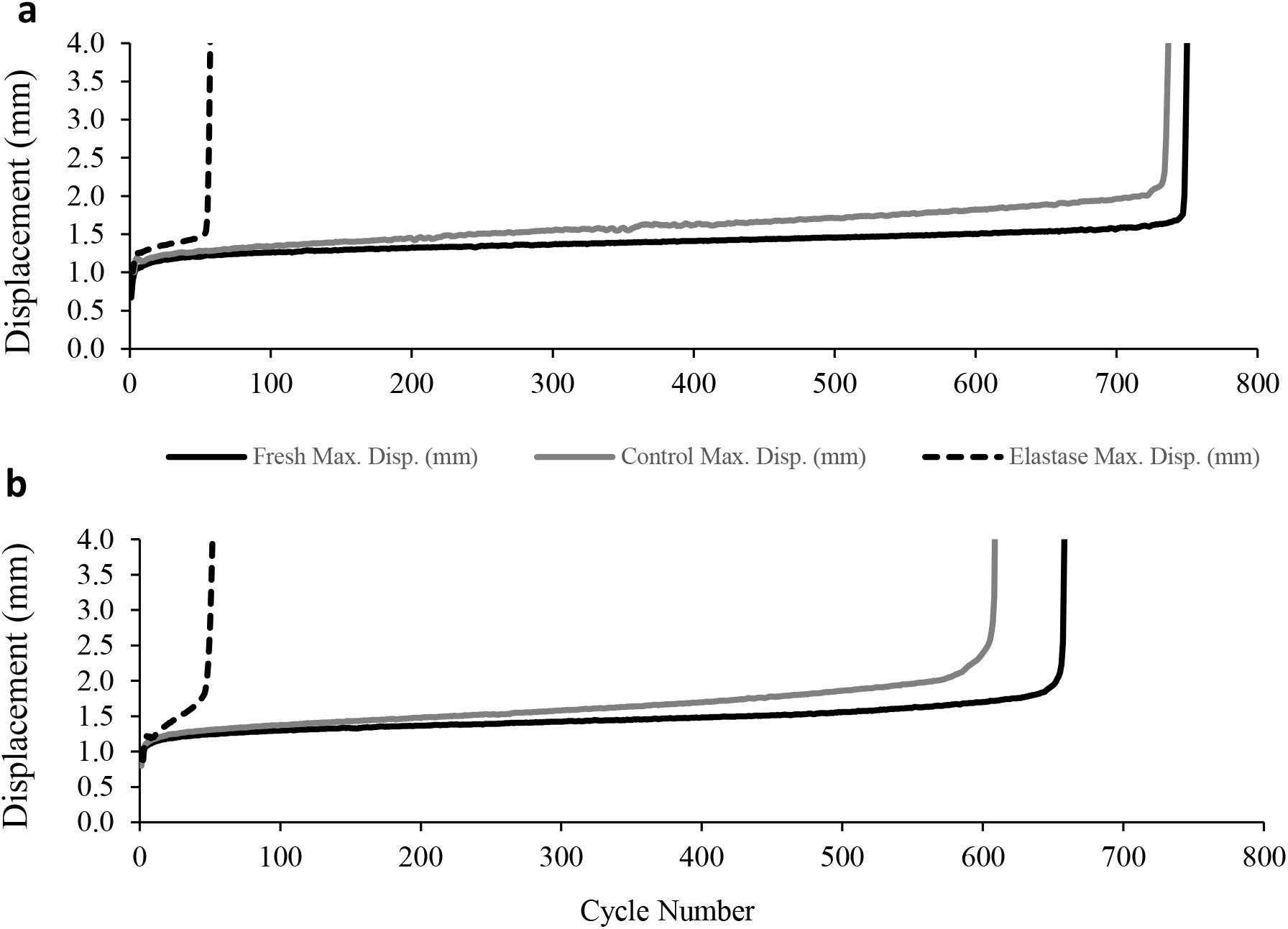
Typical creep curves of young SDFT IFM (a) and old SDFT IFM (b). The maximum displacement at each loading cycle during fatigue testing of Fresh (black line), Control (grey line) and Elastase treated (dotted line) samples are shown.

**Figure 4.**
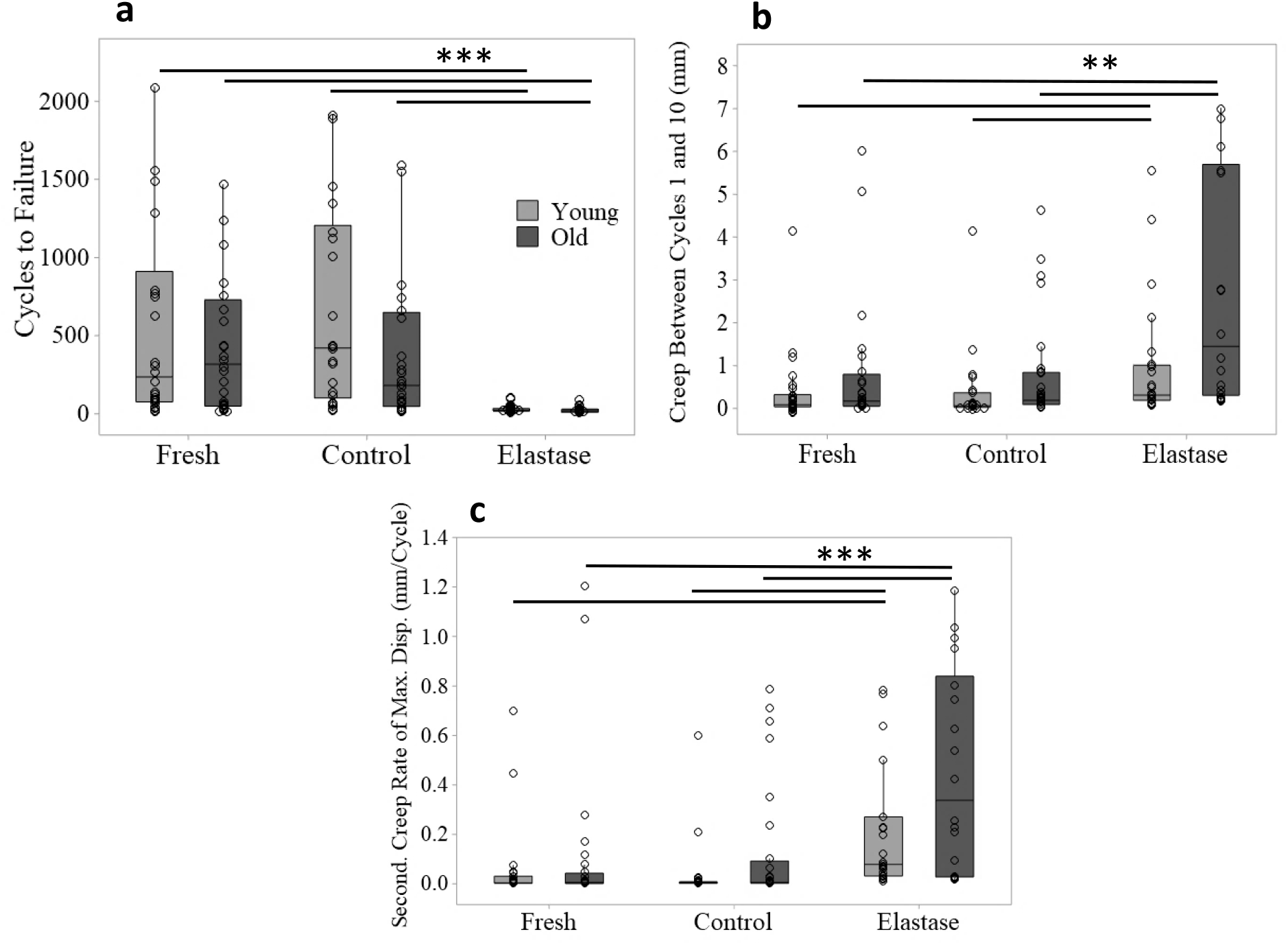
Effect of elastin depletion on IFM fatigue properties in young and old samples. Data compares cycles to failure (a), creep between cycles 1 and 10 (b) and secondary creep rate of maximum displacement (c) for young and old SDFT IFM samples from fresh, control and elastase treated groups (n=5/age group; total samples tested: 15/treatment/tendon). Significant differences are flagged with: **p<0.01 and ***p<0.001 (Cycles to Failure: normally distributed – ANOVA, all remaining IFM fatigue data not normally distributed – Mann-Whitney test).

Data show a significant reduction in the number of cycles to failure in elastase treated samples compared to both fresh and control groups (Figure 3, Figure 4a). Results also show that elastase treatment led to a significant increase in creep between cycles 1 and 10 (Fig. 4b) and secondary creep rate, compared to both fresh and control groups (Figure 4c). The response to elastase treatment did not differ significantly between young and old samples, however there was notably greater variability in aged samples, potentially masking any age- related changes (Figure 4).

### IFM Recovery Properties

The analysis of IFM recovery images is shown in Fig 5. Data demonstrated a significant increase in the angular deviation at the peak of applied load in the elastase group compared to both fresh and control groups (Figure 5a), indicating increased fascicle sliding after elastase treatment. On removal of load, the percentage recovery (figure 5b) was significantly lower in the elastase group compared to either of the other groups, and remained significantly reduced even after a hold period (Figure 5c).

**Figure 5.**
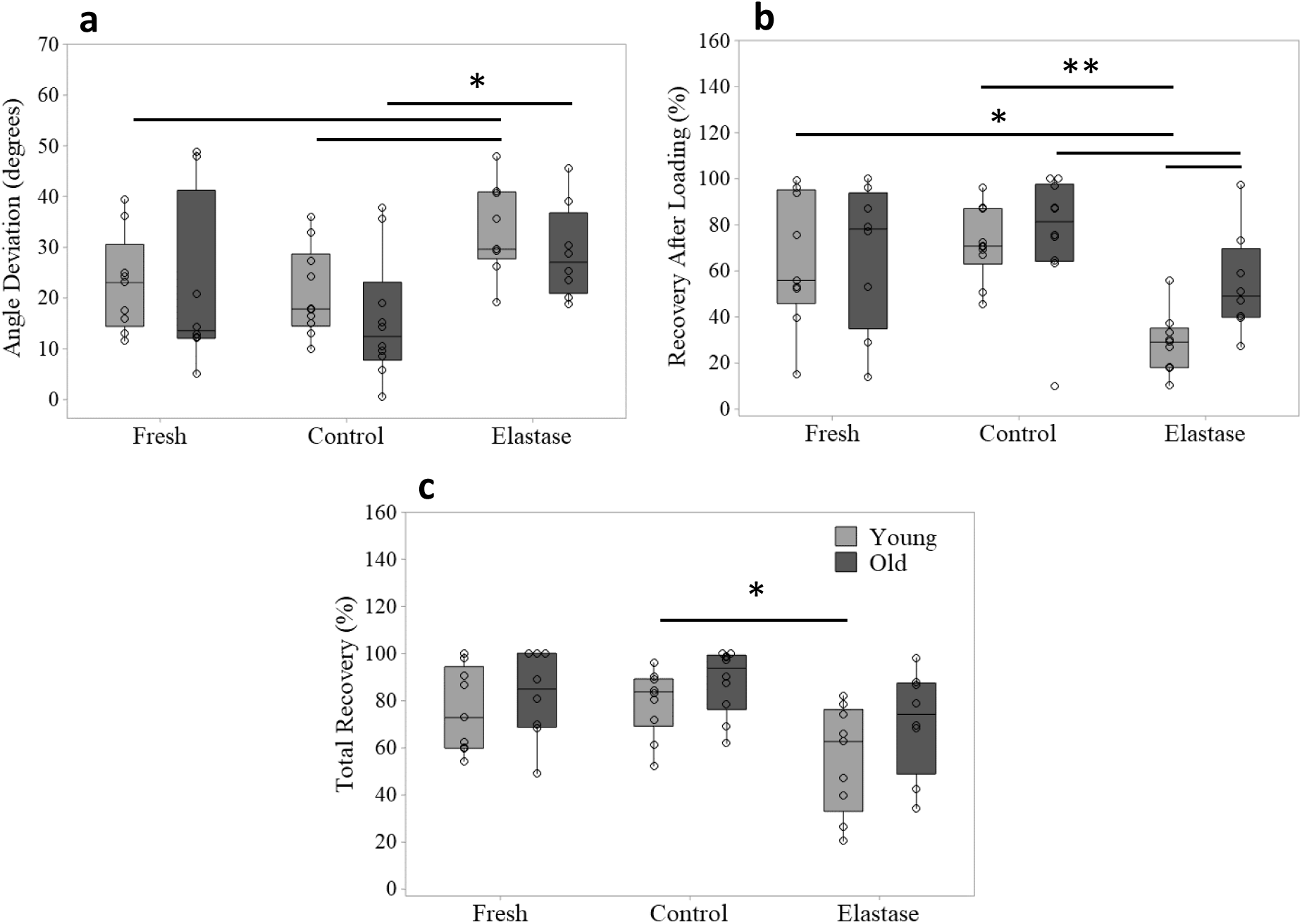
Effect of elastin depletion on IFM loading response and recovery. Loading and recovery were visually monitored by tracking markers across the IFM samples. Angle deviation (degrees) of lines was first determined under the application of 75% of the predicted failure extension (a). Load was removed, and then recovery of lines relative to their start point was measured immediately after loading (b) and 10 seconds after the removal of load (c). Graphs compare young and old SDFT IFM samples in the fresh, control and elastase treated groups (n=5/age group; total samples tested: 2/treatment/tendon). Significant differences are flagged with: *p<0.05 and **p<0.01 (data non-normally distributed – Mann-Whitney test). For more details regarding loading protocol refer to Figure 1.

In aged samples, overall trends were similar, but it was notable that immediate recovery after loading was significantly better in the aged then young elastase treatment group (Figure 5b). There were no significant differences between fresh and control groups in any of the calculated parameters.

## Discussion

This is the first study to elucidate the regional-specific influence of elastin on energy storing tendon mechanics, exploring the impact of elastin depletion on the quasi-static and viscoelastic behaviour of the IFM and fascicles in the context of ageing. Data support the hypothesis that elastin depletion exclusively impacts IFM mechanical properties, and also illustrate a consistent trend towards a more pronounced effect in young samples. All elastase-treated IFM samples consistently show a significantly reduced ability to withstand applied load, resist fatigue loading, and recover after loading.

This study specifically selected the equine model, owing to its relevance as an energy storing tendon with similar structure, disease pathology and epidemiology to that seen in human energy storing tendons [14]. This is essential for the current study, in which we hypothesise the importance of specific regions of tendon (the IFM), the structure and composition of which is similar between species [25].

The need to adopt a large animal model requires the use of enzymatic digestion approaches to carry out structure-function studies, which is associated with a number of limitations [15, 26]. Enzymatic digestion studies necessitate incubation within a buffer solution, with previous work demonstrating that the buffer solution alone can cause swelling and likely impact tendon mechanics [27, 28]. Appropriate buffer-only controls were utilised to help differentiate the impact of buffer solutions and the enzymes, and these demonstrated that mechanical changes were specific to the digest group. However, it is also well acknowledged that enzyme efficacy and specificity must be considered for targeted digestion assays. Use of proteinase-free enzyme preparations and inclusion of a trypsin inhibitor to prevent collagen degradation [18] provided the ability to remove 70% of elastin from tendon without disrupting the collagen matrix. However, elastase treatment did cause significant GAG depletion (>80%).

Some previous studies have reported a similar reduction of GAG content in other tissues exposed to elastase, likely occurring because GAG is less tightly bound within the extracellular matrix than other matrix components, and thus easily released by any disruption [15, 29, 30]. Proteoglycans interact with collagens and elastin microfibrils via their GAG sidechains, where they are believed to contribute to microfibril integration into the extracellular matrix [31]. Therefore, we performed additional experiments to determine if GAG removal alone affected IFM and fascicle mechanical properties. The results demonstrated no discernible effects of 90% GAG removal on IFM or fascicle mechanical behaviour (Supplementary Information). These findings support previous studies showing that tendon mechanical properties are not significantly affected by GAG digestion [32, 33] and provide confidence that any changes in tissue mechanics originate from loss of elastin and not GAGs.

Data demonstrate no changes to fascicle quasi-static or viscoelastic properties post-elastase treatment. With the majority of elastin in tendon localising to the IFM, and fascicles comprising more than 90% collagen, these findings are perhaps not surprising [9]. By contrast, after elastase treatment, IFM failure load and stiffness were both significantly reduced compared to both fresh and control groups, and data demonstrated a significant increase in hysteresis in elastin depleted IFM samples, and such a significant increase in the levels of primary and secondary creep, and reduction in fatigue resistance, that most samples failed immediately. Unexpectedly, ageing had little effect on IFM response to elastase treatment, with only small decreases in failure extension and initial hysteresis evident when compared to young elastase treated samples. Indeed, we did not identify any decrease in fatigue resistance with ageing in the control samples. This is in contrast to previous studies which identified a significant decrease in IFM fatigue resistance with ageing [11]. The reasons for this remain unclear, but may be a result of variability in the age-associated changes in elastin (reduced total amount, increased fragmentation and increased crosslinking).

Interestingly, no changes were observed in the failure extension of SDFT IFM samples after elastase treatment. With increased hysteresis evident in these samples, this finding was surprising, but we speculate that it arises from the protocol adopted after the preconditioning. Sample extension was normalised after the 10 preconditioning cycles and prior to the pull to failure as per previous studies [5], meaning any irrecoverable extension that occurred during the preconditioning cycles was not included in the reported values. With our results also demonstrating poor recoverability of the IFM in elastase treated samples, and previous studies showing increased lengthening of elastase treated ligaments under pre-stress [18] it seems likely that the elastase treated IFM extended notably during those initial cycles, as the IFM was less able to sustain applied load.

Taken together, our data indicate that elastase treatment specifically impacts the IFM region of tendon without affecting fascicle mechanical behaviour, and elastin depletion leads to the IFM becoming less able to withstand load, with reduced fatigue resistance. Few previous studies have investigated the effect of elastase treatment on fascicles specifically, although it has been shown that elastin depletion of rat tail tendon fascicles resulted in a reduction in failure stress and strain [15]. It is unclear why these findings are in contrast to those we report here, although significantly longer digestion times and higher incubation temperatures were used by Grant et al [15]. No previous studies have assessed the influence of elastin removal on IFM mechanics specifically. A number of studies have considered the impact of elastin depletion on whole tendon or ligament mechanics, demonstrating a reduction in tissue stiffness and/or failure stress post-elastase digestion [18, 34]. However, the effects of elastase on whole tendon viscoelastic properties remain unclear, with some studies reporting increased hysteresis [34], whilst others show no alterations [18, 35]. These differences are likely a combination of inherent differences between tendon and ligament response to elastase treatment, as well as differences in elastase treatment and mechanical testing protocols used. To the authors’ knowledge, no previous studies have directly investigated the effect of elastin depletion on tissue fatigue properties.

IFM testing in the current study was carried out in shear, pulling the opposing ends of adjacent fascicles, and demonstrating that the IFM possessed reduced ability to resist shear stresses after elastase treatment. Interestingly, whilst no comparable IFM mechanical tests exist, there are a number of studies investigating the effect of elastase treatment on either the transverse or shear mechanical properties of tendon, which would likely augment the influence of the IFM on resulting data. These studies report decreased shear and transverse stresses in both tendon and ligament after elastin depletion [17, 36]. Taken together, these results support the hypothesis that elastin provides a mechanical link between fascicles, providing the capacity to resist shearing.

Direct optical measurement of local strains in the IFM during loading and recovery provided greater insight into the mechanical behaviour reported. Data revealed a significant increase in the angular deviation during loading in the elastase group compared to both fresh and control groups, suggesting that elastase treated IFM samples stretched and sheared more than fresh and control samples, when subjected to similar displacements. The additional IFM extensibility in elastase treated samples appears contradictory to the earlier reported lack of change in IFM failure extension. However, IFM failure extension data only reported extension in the final pull to failure test, and not that occurring during preconditioning cycles. Angular deviation measurements will determine the sum of all extension from the beginning of the first loading cycle, and imaging demonstrated that the IFM responded immediately to applied load with irrecoverable extension and shear.

Optical imaging also enabled a closer investigation of recovery behaviour, showing a significant decrease in the percentage recovery of IFM sliding after load was removed in the elastase group compared to all other treatment groups. Indeed, while recovery in fresh and control samples was close to 70%, this reduced to 30% in elastase treated samples from young donors. Providing a hold period in the unloaded state for further recovery resulted in little further change in fresh and control samples, probably because most of the recovery had already been observed immediately after load was removed. By contrast, it was interesting to note that elastase treated samples from young donors showed continuing recovery during the holding period. However, the total recovery of these samples was still significantly lower than that seen in control groups.

Unexpectedly, immediate recovery from loading in elastase treated samples was significantly better in old than young IFM samples. It has previously been established that tendon elastin content decreases with ageing [9], from which it can be inferred that elastase will have less impact on aged samples. However, these findings do suggest that age-associated changes in IFM structure may lead to alternative matrix components contributing to IFM recoil in older tendons.

Taken together, data imply that elastase treatment results in increased IFM sliding and reduced ability of the IFM to withstand loading or recoil after the load is removed, all contributing to the observed reduction in fatigue resistance.

## Conclusions

These findings are of crucial importance to structure-function studies, allowing a new level of insight into the hierarchical mechanics of tendon and highlighting the important contribution of elastin to tendon mechanical properties. Data demonstrate that maintenance of a functional elastin network within the IFM is critical to maintain IFM and, consequently, tendon integrity.

## Acknowledgements

The authors would like to thank Dr Stephen Thorpe for his assistance with the statistical analysis and Dr Tony Cheung for creating the Matlab code for image analysis.

## Data availability statement

All data are available from the authors on reasonable request

## References

[1] Kannus, P., Structure of the tendon connective tissue, Scandinavian journal of medicine & science in sports 10 (2000) 312–20.

[2] Kastelic, J., Galeski, A., Baer, E., The multicomposite structure of tendon, Connective tissue research 6 (1978) 11–23.

[3] Biewener, A.A., Muscle-tendon stresses and elastic energy storage during locomotion in the horse, Comparative Biochemistry and Physiology Part B: Biochemistry and Molecular Biology 120 (1998) 73–87.

[4] Minetti, A.E., Ardig, O.L., Reinach, E., Saibene, F., The relationship between mechanical work and energy expenditure of locomotion in horses, J Exp Biol 202 (1999) 2329–38.

[5] Thorpe, C.T., Godinho, M.S., Riley, G.P., Birch, H.L., Clegg, P.D., Screen, H.R., The interfascicular matrix enables fascicle sliding and recovery in tendon, and behaves more elastically in energy storing tendons, Journal of the Mechanical Behavior of Biomedical Materials 52 (2015) 85–94.

[6] Thorpe, C., Udeze, C.P., Birch, H.L., Clegg, P.D., Screen, H.R.C., Specialization of tendon mechanical properties results from interfascicular differences, Journal of the Royal Society, Interface / the Royal Society 9 (2012) 3108–17.

[7] Thorpe, C.T., Peffers, M.J., Simpson, D., Halliwell, E., Screen, H.R., Clegg, P.D., Anatomical heterogeneity of tendon: Fascicular and interfascicular tendon compartments have distinct proteomic composition, Scientific Reports 6 (2016) 20455.

[8] Thorpe, C.T., Karunaseelan, K.J., Ng Chieng Hin, J., Riley, G.P., Birch, H.L., Clegg, P.D., Screen, H.R.C., Distribution of proteins within different compartments of tendon varies according to tendon type, Journal of Anatomy 229 (2016) 450–458.

[9] Godinho, M.S.C., Thorpe, C.T., Greenwald, S.E., Screen, H.R.C., Elastin is Localised to the Interfascicular Matrix of Energy Storing Tendons and Becomes Increasingly Disorganised With Ageing, Scientific Reports 7 (2017) 9713.

[10] Thorpe, C.T., Udeze, C.P., Birch, H.L., Clegg, P.D., Screen, H.R., Capacity for sliding between tendon fascicles decreases with ageing in injury prone equine tendons: a possible mechanism for age-related tendinopathy?, Eur Cell Mater 25 (2013) 48–60.

[11] Thorpe, C.T., Riley, G.P., Birch, H.L., Clegg, P.D., Screen, H.R.C., Fascicles and the interfascicular matrix show decreased fatigue life with ageing in energy storing tendons, Acta Biomater 56 (2017) 58–64.

[12] Grant, T.M., Thompson, M.S., Urban, J., Yu, J., Elastic fibres are broadly distributed in tendon and highly localized around tenocytes, Journal of Anatomy 222 (2013) 573–579.

[13] Thorpe, C.T., Clegg, P.D., Birch, H.L., A review of tendon injury: why is the equine superficial digital flexor tendon most at risk?, Equine Vet J 42 (2010) 174–80.

[14] Patterson-Kane, J.C., Rich, T., Achilles tendon injuries in elite athletes: lessons in pathophysiology from their equine counterparts, Ilar j 55 (2014) 86–99.

[15] Grant, T.M., Yapp, C., Chen, Q., Czernuszka, J.T., Thompson, M.S., The Mechanical, Structural, and Compositional Changes of Tendon Exposed to Elastase, Annals of Biomedical Engineering 43 (2015) 2477–2486.

[16] Eekhoff, J.D., Fang, F., Kahan, L.G., Espinosa, G., Cocciolone, A.J., Wagenseil, J.E., Mecham, R.P., Lake, S.P., Functionally Distinct Tendons From Elastin Haploinsufficient Mice Exhibit Mild Stiffening and Tendon-Specific Structural Alteration, J Biomech Eng 139 (2017) 1110031–9.

[17] Fang, F., Lake, S.P., Multiscale mechanical integrity of human supraspinatus tendon in shear after elastin depletion, J Mech Behav Biomed Mater 63 (2016) 443–455.

[18] Henninger, H.B., Underwood, C.J., Romney, S.J., Davis, G.L., Weiss, J.A., Effect of elastin digestion on the quasi-static tensile response of medial collateral ligament, J Orthop Res 31 (2013) 1226–33.

[19] Farndale, R.W., Buttle, D.J., Barrett, A.J., Improved quantitation and discrimination of sulphated glycosaminoglycans by use of dimethylmethylene blue, Biochim Biophys Acta 883 (1986) 173–7.

[20] Birch, H.L., Bailey, A.J., Goodship, A.E., Macroscopic ‘degeneration’ of equine superficial digital flexor tendon is accompanied by a change in extracellular matrix composition, Equine Vet J 30 (1998) 534–9.

[21] Legerlotz, K., Riley, G.P., Screen, H.R.C., Specimen dimensions influence the measurement of material properties in tendon fascicles, Journal of biomechanics 43 (2010) 2274–80.

[22] Thorpe, C.T., Udeze, C.P., Birch, H.L., Clegg, P.D., Screen, H.R., Specialization of tendon mechanical properties results from interfascicular differences, J R Soc Interface 9 (2012) 3108–17.

[23] Thorpe, C.T., Riley, G.P., Birch, H.L., Clegg, P.D., Screen, H.R., Fascicles and the interfascicular matrix show adaptation for fatigue resistance in energy storing tendons, Acta Biomater 42 (2016) 308–15.

[24] Screen, H.R.C., Bader, D.L., Lee, D.a., Shelton, J.C., Local strain measurement within tendon, Strain 40 (2004) 157–163.

[25] Patel, D., Spiesz, E.M., Thorpe, C.T., Birch, H.L., Riley, G.P., Clegg, P.D., Screen, H.R., Energy storing and positional human tendons: mechanics and changes with ageing, Int J Exp Path 97 (2016) A3–A3.

[26] Beenakker, J.-W.M., Ashcroft, B.a., Lindeman, J.H.N., Oosterkamp, T.H., Mechanical properties of the extracellular matrix of the aorta studied by enzymatic treatments, Biophysical journal 102 (2012) 1731–7.

[27] Screen, H.R., Chhaya, V.H., Greenwald, S.E., Bader, D.L., Lee, D.A., Shelton, J.C., The influence of swelling and matrix degradation on the microstructural integrity of tendon, Acta Biomater 2 (2006) 505–13.

[28] Safa, B.N., Meadows, K.D., Szczesny, S.E., Elliott, D.M., Exposure to buffer solution alters tendon hydration and mechanics, Journal of Biomechanics 61 (2017) 18–25.

[29] Smith, L.J., Byers, S., Costi, J.J., Fazzalari, N.L., Elastic fibers enhance the mechanical integrity of the human lumbar anulus fibrosus in the radial direction, Annals of Biomedical Engineering 36 (2008) 214–223.

[30] Jacobs, N.T., Smith, L.J., Han, W.M., Morelli, J., Yoder, J.H., Elliott, D.M., Effect of orientation and targeted extracellular matrix degradation on the shear mechanical properties of the annulus fibrosus, Journal of the Mechanical Behavior of Biomedical Materials 4 (2011) 1611–1619.

[31] Kielty, C.M., Sherratt, M.J., Shuttleworth, C.A., Elastic fibres, Journal of Cell Science 115 (2002) 2817–2828.

[32] Legerlotz, K., Riley, G.P., Screen, H.R.C., GAG depletion increases the stress-relaxation response of tendon fascicles, but does not influence recovery, Acta Biomaterialia 9 (2013) 6860–6866.

[33] Lujan, T.J., Underwood, C.J., Jacobs, N.T., Weiss, J.a., Contribution of glycosaminoglycans to viscoelastic tensile behavior of human ligament, Journal of Applied Physiology 106 (2009) 423–431.

[34] Millesi, H., Reihsner, R., Hamilton, G., Mallinger, R., Menzel, E.J., Biomechanical properties of normal tendons, normal palmar aponeuroses and palmar aponeuroses from patients with dupuytren’s disease subjected to elastase and chondroitinase treatment, Connective Tissue Research 31 (1995) 109–115.

[35] Svärd, A., Hammerman, M., Eliasson, P., Elastin levels are higher in healing tendons than in intact tendons and influence tissue compliance, bioRxiv (2020) 2020.05.26.065433.

[36] Henninger, H.B., Valdez, W.R., Scott, S.A., Weiss, J.A., Elastin governs the mechanical response of medial collateral ligament under shear and transverse tensile loading, Acta Biomater 25 (2015) 304–12.

