## Supplementary Information for "Elastase treatment of tendon specifically impacts the mechanical properties of the interfascicular matrix"

### Effect of glycosaminoglycan depletion on fascicle and IFM mechanical properties

#### Methods

##### To assess the effect of glycosaminoglycan (GAG) depletion on fascicle and IFM mechanical properties, fascicles and IFM samples were dissected from the SDFT from young horses (n = 3; aged 4 – 5 years) as described in the main methods. GAG content was depleted by digestion with 0.5U/ml Chondroitinase ABC (ChABC, AMS Biotechnology; E1028-10) in Tris buffer (50mM Tris, 60mM sodium acetate, 0.02% BSA, pH8) overnight at 37 °C with mixing (n=4-6/tendon). Two control groups were included; fresh controls were subjected to mechanical testing within 2 hours of dissection (n=4-6/tendon) and buffer controls were incubated overnight at 37 °C in Tris buffer in the absence of ChABC (n=4-6/tendon). Post-incubation, samples were rinsed with DMEM, and stored on DMEM-dampened tissue paper prior to mechanical testing. Fascicle and IFM quasi-static mechanical properties were determined following methods described in the main paper.

##### To determine the efficacy of ChABC on GAG removal, GAG content was measured post-mechanical testing. Groups of 3 samples from each experimental condition were combined to provide sufficient tissue to accurately measure weight. Samples were freeze-dried overnight, prior to the determination of dry weight. Samples were papain digested overnight at 60 °C, and GAG content determined in triplicate using the DMMB assay [19]. GAG content is displayed as µg/mg tissue dry weight.

##### Statistical significance was tested using nested one-way ANOVA with Tukey’s multiple comparisons.

#### Results

##### ChABC treatment resulted in a reduction in GAG content by 87.28 ± 2.46% compared to fresh controls (Fig. S1a). GAG depletion had no effect on fascicle or IFM viscoelastic or quasi-static properties, when compared to fresh or buffer control samples (Fig. S1b-i).

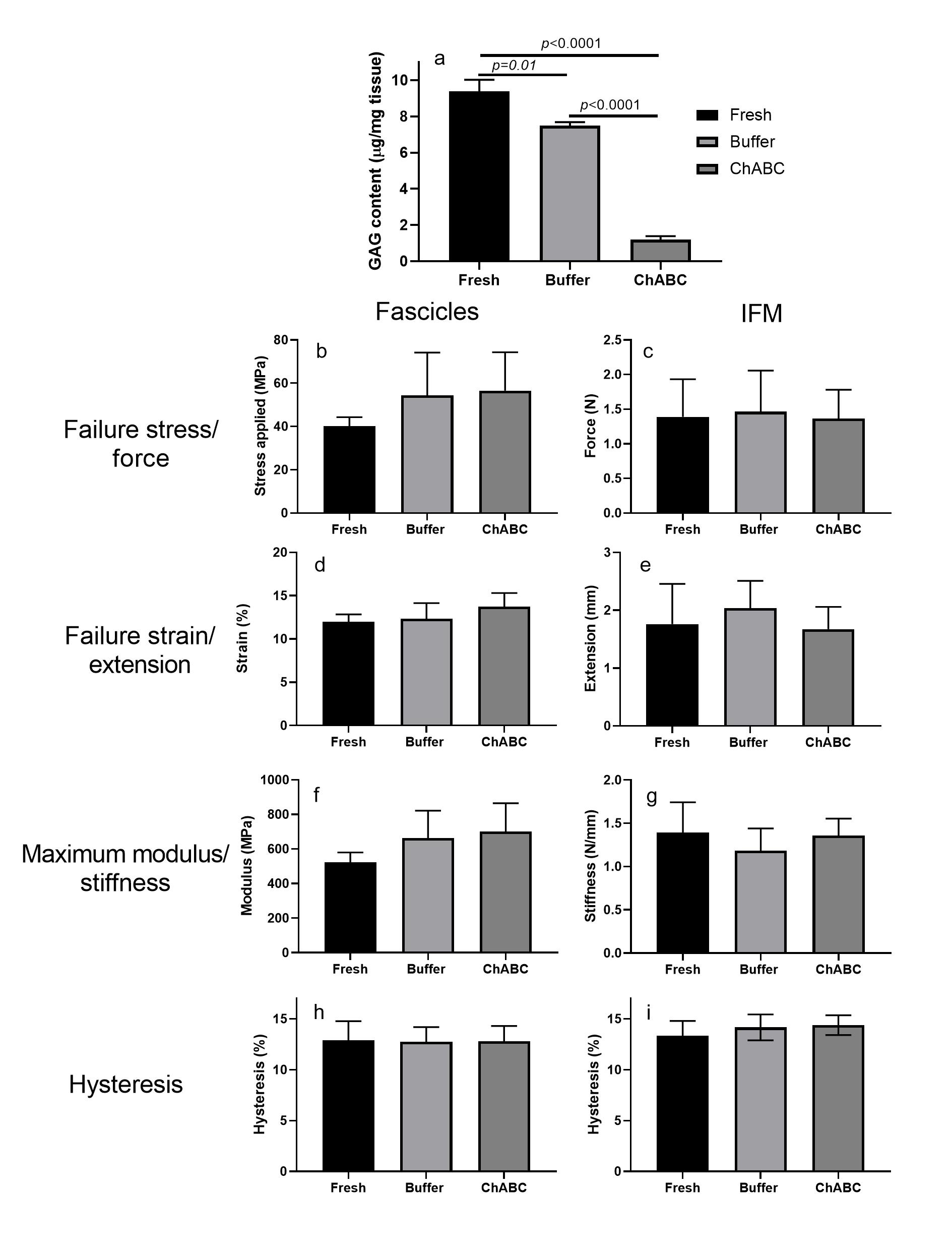

##### Figure S1. GAG depletion does not affect fascicle or IFM mechanical properties. Chondrotinase ABC treatment resulted in a significant decrease in GAG content compared to fresh and buffer controls (a). Fascicle and IFM failure properties (b-e) and maximum modulus/stiffness (f,g) showed no significant differences between groups. Hysteresis (h,i) also did not differ between control and treated groups.
